# A comparison of cognitive-motor integration performance and resting state functional brain network connectivity in female athletes suggests intact motor network and visuomotor skill in those with a concussion history

**DOI:** 10.1101/2020.10.21.348425

**Authors:** Alanna E Pierias, Diana E Gorbet, Magdalena Wojtowicz, Andrea Prieur, Lauren Sergio

## Abstract

Structural neural changes following concussion are often not captured by standard imaging techniques. However, there is growing evidence that damage to white matter tracts and change in functional network connectivity may be observed following concussive injury. We investigated behavioural performance on a cognitive-motor integration (CMI) task in conjunction with alterations in resting state functional connectivity (rs-FC) in brain networks in a population of 30 female varsity athletes, with 16 having a previous history of concussion. Behavioural performance on accuracy, timing, and trajectory measures of a CMI task were assessed between the concussion history group and the control group. Rs-FC within the nodes of the Default Mode Network (DMN), Dorsal Attention Network (DAN), the Frontoparietal Network, and the Anterior Cerebellar Lobule Network, was assessed against performance scores on accuracy, timing, and trajectory measures. Main findings indicate no difference in behavioural performance between those with concussion history and those without, in contrast to previous findings in a group of primarily male varsity athletes. In addition, no difference in rs-FC was noted in correlation with behavioural performance scores on either accuracy, timing, or trajectory. These findings may suggest sex-related differences in performance on a CMI task, and a resiliency in both functional network connectivity and visuomotor skilled performance in varsity female athletes.

## Introduction

Concussive injury has been shown to induce an altered metabolic state in the brain following injury (1,2). The length of time for complete recovery from this state is not completely understood. The vulnerability to subsequent concussive injury during the recovery period indicates an increased danger of returning an athlete to sport too soon after a concussion, and an increased potential for second impact syndrome (1,3,4), chronic traumatic encephalopathy (5,6) and even death (1,3,7,8). Therefore, there is a need to better understand the behavioural and neurological effects of concussion in athletes and others who are highly motivated to return to activity following injury.

Generally, athletes are required to maintain a high level of cognition while executing precise motor movements in order to play their sports well and to play them safely. Movements must be made in the context of game-related rules, various aspects of attention, spatial information regarding the locations of other players, and prior knowledge of how to best accomplish a given task. While there has been a great deal of research into the effects of concussion on cognition alone (9,10), there has been considerably less research into these effects on cognition integrated with movement in the context of sport and on-field performance (11). Combining thought and action is a process known as cognitive-motor integration (CMI). CMI is essential for the completion of skilled activities that involve non-standard mappings, which are relationships used when the visual information guiding the motor task does not come from the location of object one is interacting with (12,13). These mappings rely on different neural computations compared to standard, direct mapping (the spatial location of the viewed object is the same as the motor goal) that must incorporate the spatial dissociation of gaze, attention, and overt motor output. CMI is often required for complex skills, making it crucial for work, duty, sport, and daily life. Importantly, we have observed impaired CMI in youth and adolescent athletes who had experienced a concussion but were asymptomatic (using self-report) at the time of testing (14–17). These individuals included children and adolescents who played sport at the recreational and select level (14–16), as well as adolescents who played at an elite level (17). Notably, in a previous behavioural study of varsity athletes, we observed impaired CMI performance in a sample of primarily males (16/18 participants) with a history of concussion who were asymptomatic at testing, relative to peers with no history (16). In all cases, when having to incorporate some form of cognition and non-standard mapping into their movement control during an eye-hand coordination task, there were significant behavioural performance deficits in either the timing, trajectory formation, or both relative to peers with no concussion history. These data suggest that the brain networks used for cognition and motor control are affected by concussion along a timeline longer than for tasks typically used to assess symptom resolution and readiness to resume activity.

While the post-concussion CMI studies focused on movement control impairments at a behavioural level, recent studies have investigated the specific brain areas involved in cognitive-motor integration during skilled performance in healthy young adults. Gorbet and Sergio (18) investigated brain areas involved in a non-standard mapping task where saccades and reaches were spatially dissociated through a 180° rotation of cursor feedback. In this study, the authors found that activity in the cuneus and medial premotor regions was significantly different during the task preparation phase when comparing the standard to the non-standard task. Additionally, an increase in activity in the inferior parietal lobule and cerebellum was seen during the execution of the non-standard task. In a related study, Gorbet and Sergio (19) examined brain areas involved in a non-standard mapping task where saccades and reaches were made in different spatial planes. Using functional magnetic resonance imaging (fMRI) and multi-voxel pattern analysis, the authors observed a difference in brain activation between the standard and non-standard task. Interestingly, the pattern of activation was not markedly different until closer to the motor execution phase of the task, as opposed to the initial planning stages. The areas of the brain which discriminated between the two tasks in the movement period included the medial and lateral premotor, primary motor, superior and inferior parietal lobule, middle occipital gyrus, fusiform gyrus, lingual gyrus and anterior lobe of the cerebellum.

Interestingly, a difference in CMI-related brain activation has also been shown between high-performance individuals and less-experienced individuals during rule-based skill execution (19,20), and skilled performance more generally (21). Elite performers typically show reduced neural activation relative to non-elite performers during motor performance, often in sensorimotor cortices involved in skill execution (22–25). Such reduced activity is thought to reflect “neural efficiency” developed in the control networks underlying enhanced performance. When examining the brain activity of elite performers - in this case video gamers - during an integrated cognitive-motor task, Granek et al. (20) found additional prefrontal cortex activity during the planning stages of movement, as well as a later onset of preparatory activity (19). In the context of concussion recovery in experienced performers, our group has observed that number of years of sport experience is correlated with recovery, such that those who have been playing their sport longer demonstrated faster behavioural recovery relative to less-experience athletes (26). These data suggest that sport experience mitigates concussion-related skilled behavioural impairment, putatively related to ‘neural efficiency’. The underlying brain activity controlling cognitive-motor integration in experienced athletes with a concussion history has not been studied, however, in order to directly explore this idea.

The purpose of the present study was to investigate whether there was a correlation between cognitive-motor integration performance and functional network connectivity in select athletes as a function of concussion history. Due to the sex-related differences in brain network activation during equivalent cognitive-motor skill performance in healthy individuals (27), this initial study focused on data collected from female athletes only. We hypothesize that those athletes with a history of concussion (Hx - Concussion Group) will show decreased functional network connectivity in networks required to successfully execute cognitive-motor integration compared to those with no history of concussion (NoHx - Control group). Additionally, we expect the concussion history group to show behavioural performance deficits in cognitive-motor integration when compared to controls.

## Materials and Methods

### Participants

A total of 35 participants were initially recruited through the Gorman Shore Sports injury clinic on York University Keele Campus in North York, Ontario, Canada. Out of them, 3 were excluded due to excessive head motion in the MRI scanner, and 2 were excluded due to the detection of excessive susceptibility artifacts in the images. A total of 30 participants were finally included in the study (16 with a history of concussion – asymptomatic and cleared for play). The mean age of participants was 19.8 +/− 1.5 years [TABLE 1.]

**TABLE 1.** Demographic information on the 30 female varsity athletes included in analysis, including the varsity sport they play at York University (sport), their age, the number of years they have been playing said sport (years of experience), whether they have sustained a concussion previously (Concussion History – n = no, y = yes), and how many previous concussions they have sustained throughout their life obtained through self-report (Number Previous Concussions).

| Participant Information |  |  |  |  |
| --- | --- | --- | --- | --- |
| Sport | Age | Years of Experience | Concussion History | Number Previous Concussions |
| volleyball | 21 | 9 | n | 0 |
| volleyball | 21 | 16 | n | 0 |
| hockey | 20 | 15 | n | 0 |
| soccer | 18 | 14 | n | 0 |
| soccer | 21 | 17 | n | 0 |
| basketball | 18 | 12 | n | 0 |
| hockey | 19 | 12 | n | 0 |
| hockey | 18 | 14 | n | 0 |
| hockey | 20 | 15 | n | 0 |
| hockey | 19 | 14 | n | 0 |
| hockey | 19 | 15 | n | 0 |
| hockey | 18 | 13 | n | 0 |
| hockey | 20 | 16 | y | 1 |
| soccer | 21 | 11 | y | 1 |
| hockey | 18 | 15 | y | 1 |
| hockey | 20 | 13 | y | 1 |
| volleyball | 19 | 5 | y | 1 |
| hockey | 19 | 15 | y | 1 |
| basketball | 22 | 10 | y | 1 |
| basketball | 19 | 9 | y | 2 |
| volleyball | 22 | 16 | y | 2 |
| volleyball | 19 | 13 | y | 2 |
| soccer | 21 | 15 | y | 2 |
| volleyball | 21 | 16 | y | 3 |
| basketball | 19 | 8 | y | 3 |
| soccer | 20 | 14 | y | 3 |
| hockey | 20 | 15 | y | 3 |
| volleyball | 21 | 13 | y | 4 |
| soccer | 18 | 14 | y | 5 |
| soccer | 24 | 21 | y | 6 |

All participants included in the study had no history of neurological problems, and no other injury preventing them from practicing and playing their sport. Additionally, all participants were screened using MRI safety protocols to ensure they were cleared to enter the MRI scanner. Participants were reimbursed for their time at a rate of $50 for one two-hour session. The York University Research Ethics board human participants subcommittee approved the protocol used in the experiment. The experimental protocol was also in compliance with the Declaration of Helsinki. All participants provided informed written consent prior to data collection

### Procedure

Participants completed an imaging protocol and behavioural data collection in a single session. To help ensure consistent resting state data between individuals, the imaging protocol was completed prior to behavioural data collection for all participants.

Participants were scanned in a supine position with a 3T Siemens Magnetom Trio scanner at York University. For the resting state functional magnetic resonance imaging scan (rs-fMRI) each participant was instructed to rest (but remain awake) with their eyes open, focus on a fixation cross, and not to think of anything in particular. The participants were scanned in the sagittal plane, using a 32-channel head coil.

Anatomical images were acquired using a T1-weighted MPRAGE sequence (Field of View (FOV) = 256 mm, flip angle (FA) = 9 degrees, repetition time (TR) = 2.3s, 192 slices, 1.0 x 1.0 x 1.0 mm voxel resolution). Functional images (rs-fMRI) were acquired using multi-echo echo-planar imaging (EPI) using the following parameters: 216 mm FOV, FA=83 degrees, slice thickness 1.6 mm (no gap), TR=3.0s, and echo times (TEs)=14, 30, 46 ms. One resting functional run consisting of 192 images was acquired for each participant.

For the behavioural collection, participants completed two computer-based visuomotor transformation tasks, one standard (V) and one non-standard (HR) (vision and action decoupled). Participants sat at a desk comfortably in front of a 10.1” ACER tablet (60Hz refresh rate) connected to a 15” Keytec™ touchpad allowing for a touch screen in both the horizontal and vertical planes. In both conditions, participants were instructed to slide their index finger of their preferred hand along the touch screen in order to displace a cursor from a central target to one of four peripheral targets (up, down, left, right) as quickly and as accurately as possible. Participants guided a crosshair cursor, viewed on a black background, to the yellow central (or home) target which, when reached successfully, changes to green. After 4000ms, a red peripheral target is presented and the central target disappears, which serves as the ‘Go’ signal for the participant to initiate movement. Once the participant successfully reaches the target and remains for 500ms, the peripheral target disappears, signaling the end of the trial. The next trial begins with the presentation of the central target after an inter-trial interval of 2000ms. Peripheral targets are located 55mm from the center target, and target diameters are 15mm. In order to ensure smooth movement of the finger during the task, participants wore a capacitive-touch glove on their preferred hand (Figure 1).

**Fig. 1.**
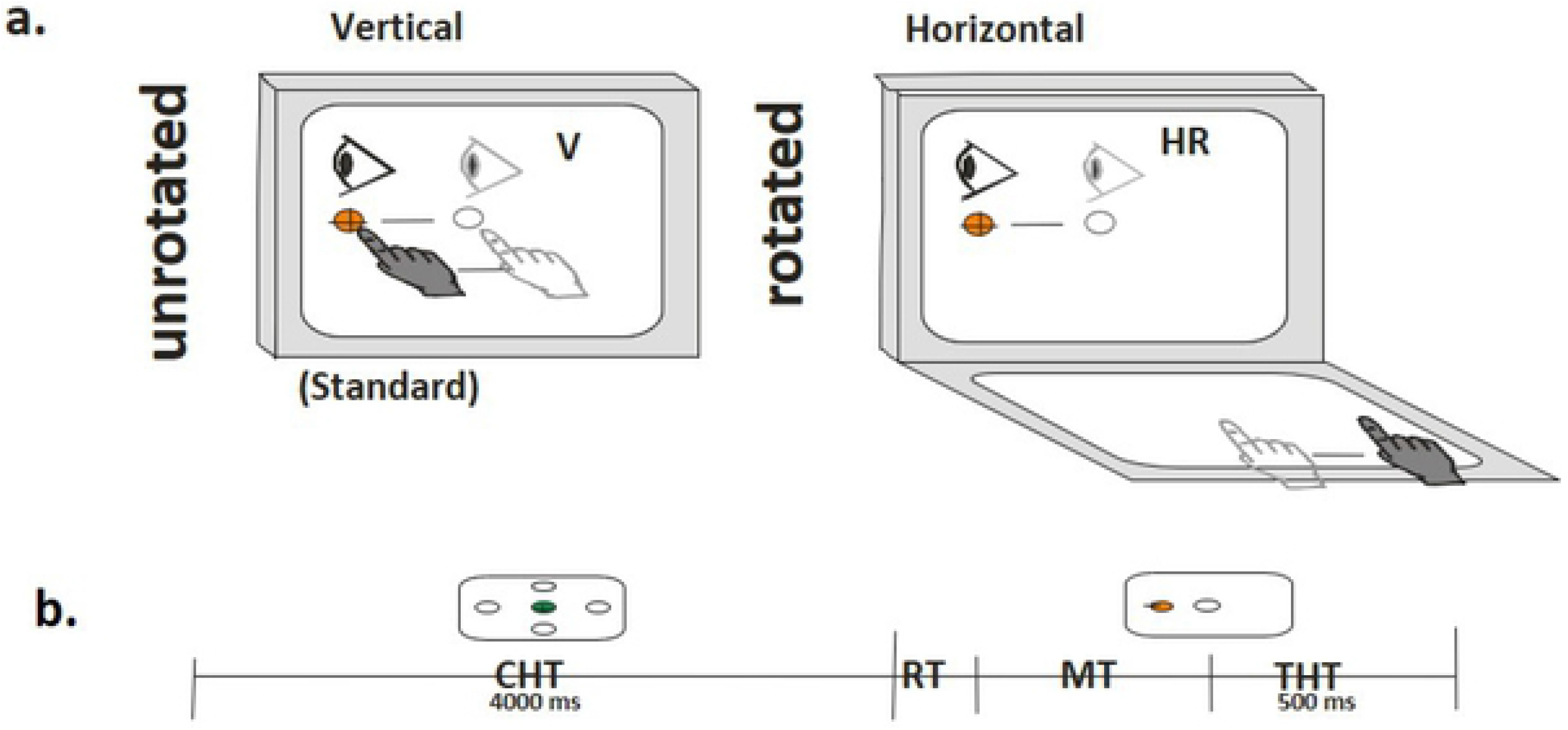
(A) Schematic drawing of both experimental conditions V (vertical, standard) and HR (Horizontal Rotated, non-standard). Visual stimuli were presented on the vertical monitor for all conditions. Green circle, light grey eye, and hand symbols denote the instructed eye and hand movements for each task. Red circles denote the peripheral (reach) target, presented randomly in one of four locations (left, up, right, and down). The dark crosshair denotes the cursor feedback provided during each condition. (B) Schematic drawing of time progression of task. centre hold time (CHT) indicates the 4000ms participants must hold in the centre target before a peripheral target appears, followed by reaction time (RT) of participant to respond to the stimulus, movement time (MT) of participant to successfully slide the cursor from the central target to the peripheral target, and target hold time (THT) signifying the 500 ms that participants must hold in the peripheral target until returning back to centre to start their next trial.

Participants completed the standard and non-standard conditions in a randomized block design. In the standard condition, participants both look and move on the vertical screen, and thereby directly interacted with the targets. Movements for the non-standard visuomotor condition are made on the horizontally oriented Keytec™ touchpad. The touchpad was calibrated such that the distance of the finger movement path on the touchpad is equivalent to the distance the cursor moves on the tablet screen. The non-standard condition includes two levels of decoupling: (i) plane change; in which participants look at the vertical screen while moving on the horizontal screen, requiring spatial recalibration, and (ii) cue reversal; in which the feedback is rotated 180° (i.e. in order to move the cursor left, you slid your finger right), requiring strategic control. Six trials in each of the four directions were completed per condition for a total of 40 trials per participant (4 directions x 5 trials x 2 conditions). Eye movements were monitored by the experimenter to ensure participants look at the correct target

### fMRI data Preprocessing

Multi-echo rs-fMRI data preprocessing was performed using the Multi-Echo Independent Components Analysis (ME-ICA) pipeline in AFNI. In order to preprocess the data using ME-ICA, dcm2nii was first used to convert the files from Dicom to NIfTI file format (28). Images were visually inspected at this stage to ensure proper conversion. Prior to preprocessing of the functional data, the first 5 volumes (acquired prior to MRI steady-state equilibrium) were eliminated. Preprocessing of the functional data included motion correction (AFNI 3dvolreg), removal of large signal transients via interpolation (AFNI 3dDespike), and slice time correction (AFNI 3dTshift). Coregistration of functional and T1-weighted anatomical images was performed (AFNI 3dAllineate) and functional data were transformed to MNI space (AFNI 3dWarp). TE-dependent denoising (ME-ICA) was implemented in AFNI as described in Kundu et al., 2013 (29). Spatial smoothing was applied to the denoised functional data using a 6-mm full-width half-maximum (FWHM) Gaussian kernel and the functional data were scaled to a mean value of 100. After preprocessing, quality control steps included verification that scans did not include excessive head motion, confirmation that functional data were accurately aligned to anatomical data, and verification that data were not excessively noisy (a minimum of 10 independent components classified as BOLD signal by ME-ICA).

### Behavioural data preprocessing

Custom-written (C++) acquisition software samples the finger’s X-Y screen position at 60Hz. Custom analysis software (Matlab, Mathworks Inc.) was used to generate a computerized velocity profile of each trial’s movement. Movement onsets and ballistic movement offsets (the initial movement prior to path corrections) were scored at 10% peak velocity, while total movement offsets were scored as the final 10% peak velocity point once the finger position is within the peripheral target. These profiles were then verified by visual inspection, and manually corrected if necessary. Trials were considered errors if the finger/cursor left the center target too early (< 4000ms), reaction time was less than 150ms or more than 8000ms, or total movement time was more than 10000ms. Trials in which the first ballistic movement exited the boundaries of the center target in the wrong direction (greater than 90° from a straight line to target) were coded as direction reversal errors, eliminated from further evaluation, and analyzed as a separate variable. The scored data was then processed to compute both movement timing and execution outcome measures.

The kinematic variables for movement timing were as follows: 1) Reaction Time (RT), the time interval (milliseconds) between the central target disappearance and movement onset; 2) Movement Time (MT), the time between movement onset and offset (millisecond), and 3) Peak Velocity (PV), the maximum velocity obtained during the ballistic movement. Kinematic variables for movement execution were: 1) Full Path Length (PLf), the total distance (resultant of the x and y trajectories) travelled between movement onset and offset (millimeters), 2) Absolute Error (AE, accuracy), the average distance from the individual ballistic movement endpoints (∑ x/n, ∑ y/n) to the actual target location (millimeters); 3) Variable Error (VE, precision), calculated as the distance between the individual ballistic movement endpoints (σ2) from their mean movement (millimeters); and 4) Percentage Direction Reversal errors (DR) were calculated as the number of trials with a deviation of greater than ±45° from the direct line between the midpoint of the central and peripheral targets compared to the number of successful trials.

#### Data Analysis

##### Behavioural

For six of the dependent variables described above (RT, MT, PV, PLf, CE, VE) raw output scores were added together in related groups to create composite scores which generated an overall performance score for Timing (RT and MT), Trajectory (PV and PLf) and Accuracy (AE and VE). In the case of the Trajectory score, PV raw output was multiplied by negative one before adding to ensure that a higher score indicated a worse performance, in line with PLf. The number of direction reversal errors (DR) was analyzed separately. To include a comparison of the performance as a function of their change from their own standard behaviour, composite scores will be composed of delta scores (standard condition subtracted from the non-standard condition).

##### Imaging

The ME-ICA denoised functional data were used in the group level analyses using FRMRIB Software Library (FSL). Specifically, Multivariate Exploratory Linear Optimized Decomposition into Independent Components (MELODIC) was used for decomposition of the functional data into independent components. Within MELODIC, a multi-session temporal concatenation approach was used. This approach concatenates the data matrices for each subject allowing MELODIC to isolate spatial patterns that are common to the group. Additionally, MELODIC uses a probabilistic ICA method that avoids over-/under-fitting with use of a Bayesian approach for estimation of the amount of Gaussian noise in the data. Therefore, arbitrary pre-specification of data dimensionality (i.e. number of components to isolate) can be avoided (30). Of the final components, we identified four a priori-selected resting state fMRI networks: the Default Mode Network (DMN), the Dorsal Attention Network (DAN), the frontoparietal network (FP), and the anterior cerebellar network (31,32) which are additionally known to be involved in CMI tasks (13,18) and commonly impacted by concussion (14,16,33). The group level results from MELODIC were input into a dual regression analysis within FSL that was utilized to back-project spatial maps and individual time series for each component and subject. This technique regresses the group ICA component maps into subject-specific temporospatial maps by applying 2 regression analyses using both temporal coherence and spatial similarity of the component (34). We used these subject-specific spatial maps to investigate the differences in functional connectivity within the chosen networks in a voxel-wise fashion between those with concussion history and controls. Assessment of group-related differences in functional connectivity within the networks of interest were performed using the Randomise function within FSL. Nonparametric permutation tests (5000 permutations) were used to detect statistically significant differences within the networks of interest. Statistical thresholding was performed using threshold-free cluster enhancement (TFCE) and a significance threshold corrected for multiple comparisons was set to p<0.05.

Using a two-group difference with continuous covariate interaction general linear model (GLM) within Randomise, we investigated whether the interaction between accuracy, timing, and trajectory performance scores, and functional network connectivity differed between the concussion history group, and the control group within resting state networks of interest. Following this, we ran a two-group difference GLM (a two-sample independent t-test) to see if there were mean differences in functional connectivity between the concussion history group and the control group.

Since the statistical models described above did not reveal statistically significant differences between groups (see Results section, below), we also tested a model with all participants collapsed into a single group to investigate whether there was an interaction between our behavioural performance scores (accuracy, timing, and trajectory composite scores modeled as covariates) and functional network connectivity across all participants.

## Results

### Behavioural

A t-test was conducted comparing performance scores of the Concussion history group to the Control group on all raw behavioural output variables (RT, MT, PV, PLf, CE, VE, DR). No significant differences were found on any of the variables. A trend towards significance was seen in PLf (p = 0.08) and PV (p = 0.09) with the Concussion history group exhibiting longer pathlengths and slower peak velocities (FIGURE 2).

**Fig. 2a & 2b.**
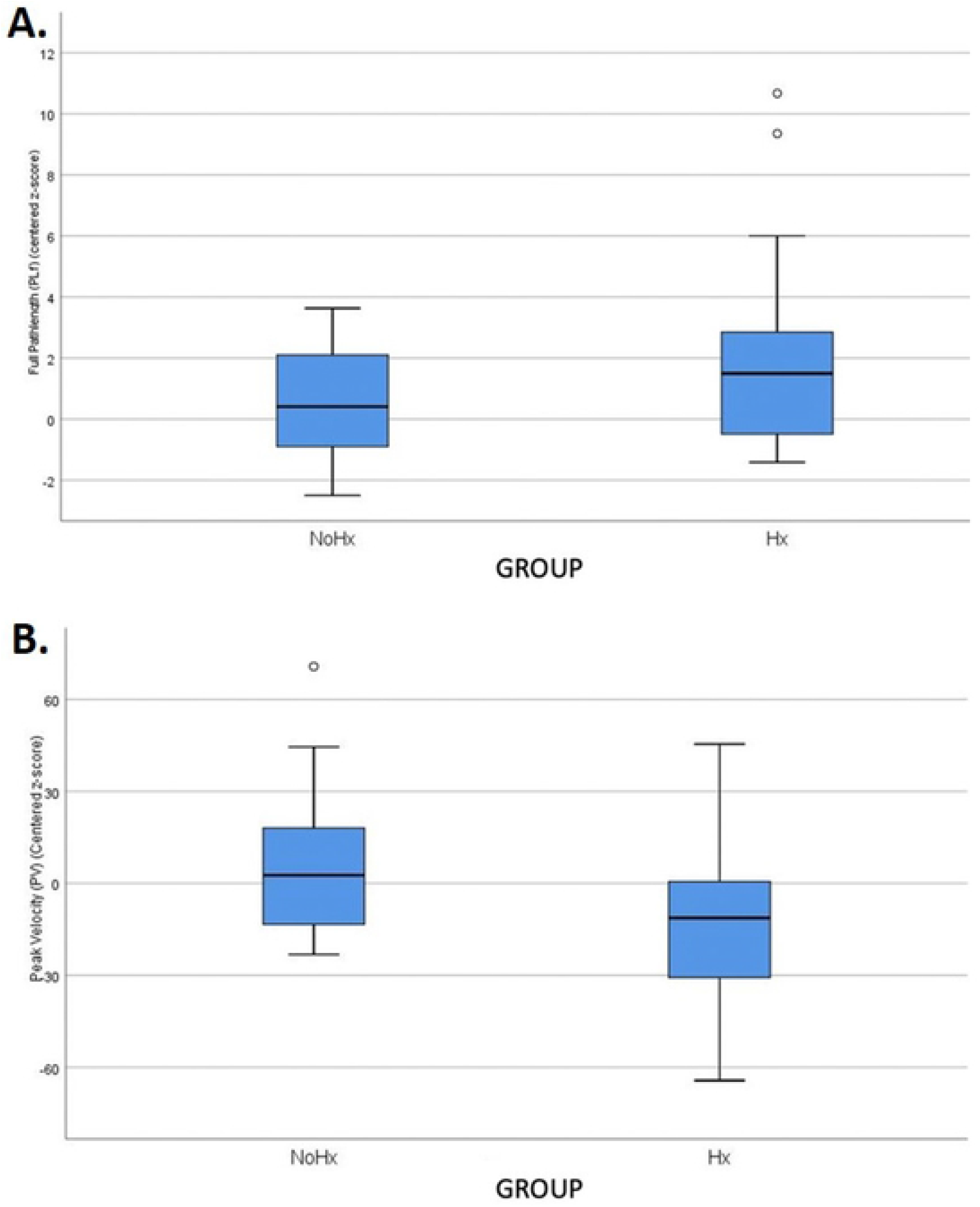
(A) Boxplot showing the centered z-scores for full pathlength kinematic outcome variable by group (NoHx – No previous concussion history, Hx – previous concussion history). (B) Boxplot showing the centered z-scores for peak velocity kinematic outcome variable by group NoHx and Hx. Error bars represent standard deviation.

A t-test was conducted comparing the Timing, Trajectory, and Accuracy composite scores of the Concussion group to the Timing, Trajectory, and Accuracy scores of the Control group. No significant differences were found in any of the three score categories (Timing, p=0.63; Trajectory, p= 0.17 ; Accuracy, p= 0.78)

A Pearson correlation was conducted comparing each composite score (Timing, Trajectory, and Accuracy) with number of previous concussions. No statistically significant differences were found for Timing (r = −0.09), Trajectory (r = 0.25) or Accuracy (r = −0.05).

Additionally, no significant difference was seen when comparing concussion history with number of direction reversal errors (p = 0.15).

### Resting state functional network connectivity

No statistically significant differences were found in the resting state functional network connectivity of the DAN, the DMN, the fronto parietal network, or the anterior cerebellar lobule network when comparing the Concussion history group to the Control group in the two-group comparison with continual covariate interaction model (p>0.05 for all; TABLE 2).

**TABLE 2.** Report of p-values for each two-group with continuous covariate interaction model for each of the four networks tested by composite score (accuracy, timing, trajectory).

| Network | Composite Score | p-value |
| --- | --- | --- |
| Default Mode Network | Accuracy | 0.73 |
|  | Timing | 0.89 |
|  | Trajectory | 0.89 |
| Dorsal Attention Network | Accuracy | 0.66 |
|  | Timing | 0.55 |
|  | Trajectory | 0.45 |
| Frontoparietal Network | Accuracy | 0.80 |
|  | Timing | 0.93 |
|  | Trajectory | 0.92 |
| Anterior Cerebellar Network | Accuracy | 0.56 |
|  | Timing | 0.52 |
|  | Trajectory | 0.71 |

No statistically significant differences were found in mean functional connectivity between the Concussion history group and the control group in the two-group unpaired t-test model for any of the resting state networks of interest (p>0.05 for all).

No statistically significant correlations were found in the DAN, the DMN, the frontoparietal network, or the anterior cerebellar lobule network for Timing, Trajectory, or Accuracy composite scores in the single-group average with additional covariate model.

Below, we include scatterplots as an illustrative example. These figures were derived by extracting the mean z-value from a node within one of our resting state networks (in this case, the Precuneus within the DMN) using the individual subject spatial maps generated by dual degression. These values were then plotted against the composite scores to illustrate the lack of a systematic relationship. (FIG 3.a b c)

**Fig. 3a, b, c.**
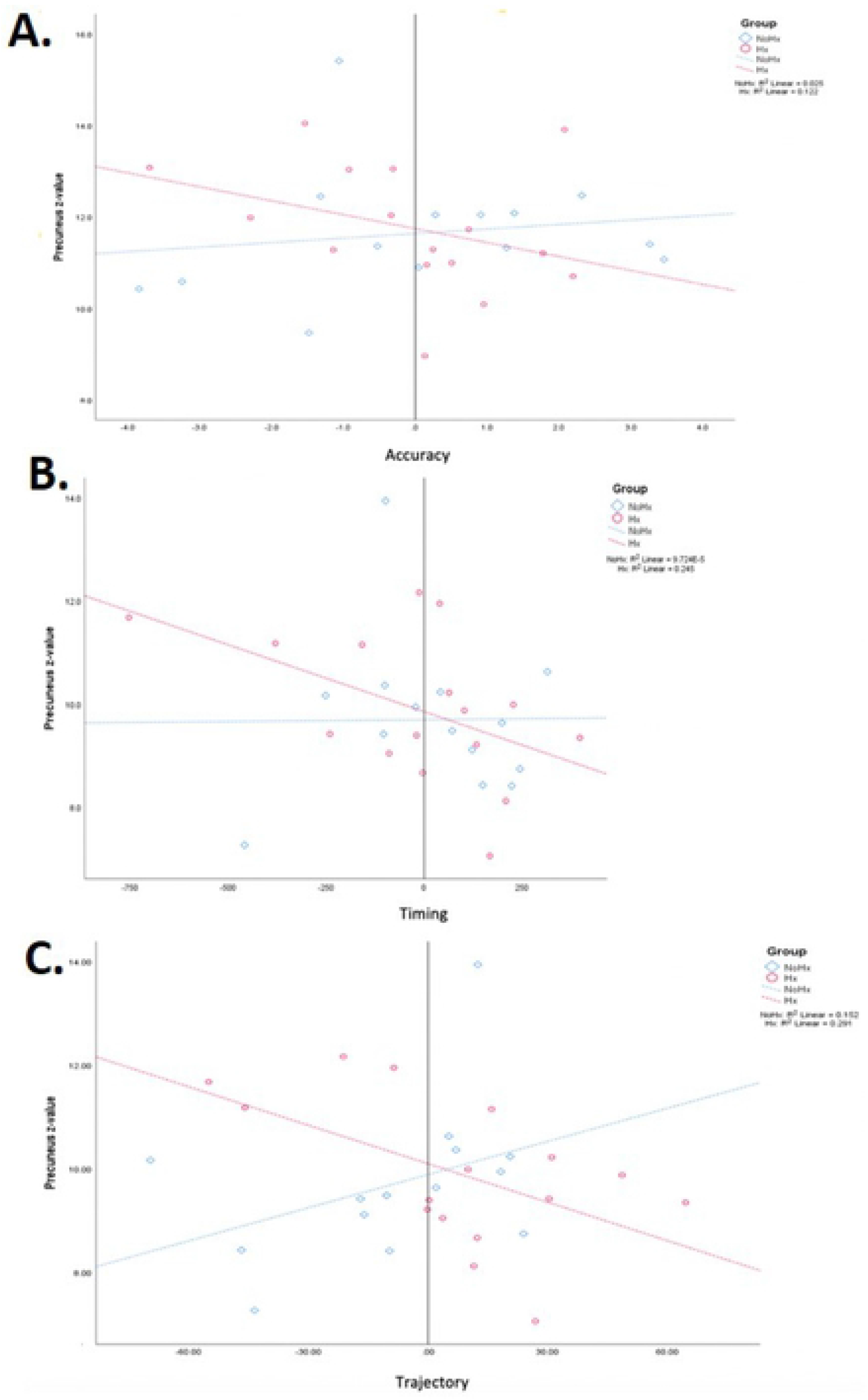
Scatterplots derived by extracting the mean z-value of the Precuneus within the DMN using the individual subject spatial maps generated by dual degression. These values were then plotted against the composite scores (A)Accuracy, (B) Timing, and (C) Trajectory. A regression line and r^2^ value is included for each group (NoHx-no previous concussion history, Hx – previous concussion history).

## Discussion

We examined the effect of concussion history on CMI performance and how this performance relates to resting-state functional network connectivity. We examined functional networks with nodes commonly impacted by concussion and that are known to be used to successfully execute CMI tasks. Contrary to our hypothesis, no differences were found in connectivity regardless of performance on our CMI task in our group of female varsity athletes. Specifically, the dorsal attention network, default mode network, frontoparietal network, and anterior cerebellar lobule network showed no increases or decreases in functional connectivity in correlation with any of our timing, trajectory, or accuracy composite scores. Additionally, having a concussion history did not impact CMI performance as measured by our composite scores and number of direction reversal errors when compared to those with no concussion history. Given previous work looking at neural connectivity impairments following concussion and mTBI which noted differences in white matter integrity (33,35) and functional network strength and activation (36–38) these results were unexpected. However, these previous studies included a mixture of mTBI and concussion patients (36), observed only males (37,38) or included males and females in the same analysis (35,36,38), and were done at the acute/subacute stage of concussion (36–38). As noted previously, the CMI task used in this experiment has shown sex-related differences in activation in the healthy brain; therefore, there is a possibility that the functional network activity following concussion is impacted differently in females than in males. Interestingly, a decrease in these differences of network activation in those with concussion history and those without has been correlated with a decrease in symptoms (38), and with time since injury (39.40). Given that all of our participants were >6 months post-injury, and deemed asymptomatic and cleared to return to play, this may also provide us with a possible explanation as to why there are no differences in functional network connectivity between the two groups.

Previous studies investigating CMI performance with and without concussion history have shown an impact of concussion on CMI performance (16,17) lasting up to two years (14). However, these previous studies were done with youth (14), included both males and females in a single group (16), or on males only (17). Given the nature of sex-related differences in cognitive performance, and brain network activation patterns in CMI tasks, there is a possibility that previously observed differences in CMI performance in those with a concussion were driven by males. Indeed, in terms of recovery from concussion, there is evidence of sex-related differences in both the rate of recovery and the form of recovery, although these findings are mixed (41,42). In humans it has been proposed that these differences may be related to neck girth (43), hormonal influences (44), and cerebral blood flow and metabolic differences (45). Additionally, recent work by Dumais et al. (46) found that males and females have a difference in activation of the default mode network, and the dorsal attention network when exposed to punishment and reward; however, no difference in activation was seen during working memory. This may provide insight into the lack of behavioural differences noted in this study and perhaps suggest that a correlation of behavioural performance and functional network activation may still exist in males. Rodent studies support these ideas. Intriguingly, a recent study by McCorkle et al. (47) observed sex-related differences in concussion recovery, whereby male mice experienced longer recovery of visuomotor skills, while female mice experienced longer recovery of emotion-related symptoms (but not visuomotor skills). To this end, further research is needed to determine whether analogous behavioural differences in CMI performance exist in male varsity athletes, and whether they exhibit a correlation between resting state functional network connectivity and CMI performance.

The capacity of the brain to continue to perform without noticeable impairment until damage reaches a critical level has been termed neuronal reserve (48–50). The athletes examined in this study all have extensive experience in their sport and many have been playing for most of their lives. While reserve is traditionally a concept applied to forms of cognition, we propose that the lack of difference observed in resting state functional network connectivity in this study may also be due to this phenomenon. Previous research observing Parkinson’s and Alzheimer’s patients suggested an active cognitive reserve, in which the normal task-related networks are recruited to a greater extent than normal to preserve performance levels (48,51). Additionally, the increased density and coherency of neurites noted by Churchill et al. (35) which strengthened with time since injury has been suggested to reflect an adaptive response to concussion, as opposed to a pathology of the injury. In the case of the current study, CMI performance scores were not reflected in resting state functional network connectivity; however, it is possible that an increased activation of these networks during execution of the task is required in those with a history of concussion in order to produce the lack of behavioural differences in performance observed.

Similarly, Hurtubise et al. (17) suggested a compensatory mechanism in elite athletes derived from extensive training of these networks in the context of their sport which allow them to perform to an acceptable baseline standard, despite there still being changes in the brain following concussion. While these authors noted a difference in performance between elite level athletes, and varsity level athletes, the study consisted of only males. The female participants we tested in the current study were varsity level athletes who have been training in their sport for an average of 15 ± 3 years. They were all asymptomatic, free of neurological disorders, and actively training in their sport at the time of testing. Therefore, it is possible that the lack of behavioural differences in performance on a CMI task between females with and without a history of concussion is derived from increased strengthening of the brain networks required to successfully perform a CMI task through extensive training in their sport. This hypothesis could be tested further by examining behavioural and resting-state functional connectivity correlates in a group of male athletes with and without concussion history. If resiliency is a contributing factor to the lack of differences in resting-state functional connectivity between athlete groups in this study, it is important to investigate further by comparing athlete to non-athlete individuals. Such a future study would mo comprehensively examine the aspect of resiliency discussed here.

Finally, the cerebellum – responsible for motor coordination, motor learning, and spatial attention (18, 52), and often not accounted for in this field of research – also plays an essential role in both standard and non-standard reaching tasks (13,53,54). Increased cerebellar activity has been noted in non-standard compared to standard visuomotor tasks, resulting from the need for corrective movements or possibly due to a role in the actual dissociation of eye and hand (18). The lack of difference in the anterior cerebellar network between those with concussion history and those without is somewhat surprising given that balance related symptoms are often observed following concussive injury (55,56). However, the cerebellum is also known to be particularly important during corrective movement stages when sensory feedback is available (52,57,58). Therefore, it would be of interest to investigate whether there is a correlation between CMI performance and activation of the anterior cerebellar network during active CMI task fMRI. Additionally, changes in white matter integrity has been linked to a poorer CMI performance in those at risk for Alzheimer’s and dementia (59). Similarly, a correlation between performance on a CMI task and white matter integrity (as measured using fractional anisotropy) has been noted in those experiencing postconcussional syndrome and age-matched controls (60) whereby decreased white matter integrity is correlated with a poorer CMI performance. If the lack of difference in functional network activation in this study is due to a reserve creating a resiliency effect achieved via years of practicing a sport, it would be important to observe whether this resiliency remains intact with progressive aging and the natural effects of age-related cognitive slowing. If there is underlying damage to the brain which has been hidden through compensation mechanisms and reserve, will this damage be increasingly evident in the long-term, or will said resiliency provide a protective effect leading to decreased or slowed cognitive decline down the road.

Overall, this study provides a starting point for many more interesting questions surrounding the sex-related and long-term effects of concussion.

## Acknowledgements

We would like to thank Joy Williams for all of our MR image acquisition, Melissa Miljanovski for assistance with data collection, York University Faculty of Health for imaging pilot funds (MW), and operating support from an NSERC discovery grant-in-aid (LS) and VISTA (CFREF)/York University Research Chair funds (LS).

## REFERENCES

1 Musumeci G, Ravalli S, Amorini AM, Lazzarino G. Concussion in Sports. Journal of Functional Morphology and Kinesiology. 2019 Jun;4(2):37.

2 Banks RE, Domínguez DC. Sports-related concussion: neurometabolic aspects. InSeminars in speech and language 2019 Aug (Vol. 40, No. 05, pp. 333–343). Thieme Medical Publishers.

3 McCrory P, Meeuwisse WH, Aubry M, Cantu RC, Dvorak J, Echemendia RJ, Engebretsen L, Johnston KM, Kutcher JS, Raftery M, Sills A. Consensus statement on concussion in sport—the 4th International Conference on Concussion in Sport held in Zurich, November 2012. PM&R. 2013 Apr;5(4):255–79.

4 Prins ML, Hales A, Reger M, Giza CC, Hovda DA. Repeat traumatic brain injury in the juvenile rat is associated with increased axonal injury and cognitive impairments. Developmental neuroscience. 2010;32(5-6):510–8.

5 McKee AC, Stein TD, Kiernan PT, Alvarez VE. The neuropathology of chronic traumatic encephalopathy. Brain pathology. 2015 May;25(3):350–64.

6 Saffary R, Chin LS, Cantu RC. From concussion to chronic traumatic encephalopathy: A review. Journal of Clinical Sport Psychology. 2012 Dec 1;6(4):351–62.

7 Harmon KG, Clugston JR, Dec K, Hainline B, Herring S, Kane SF, Kontos AP, Leddy JJ, McCrea M, Poddar SK, Putukian M. American Medical Society for Sports Medicine position statement on concussion in sport. British journal of sports medicine. 2019 Feb 1;53(4):213–25.

8 Barkhoudarian G, Hovda DA, Giza CC. The molecular pathophysiology of concussive brain injury. Clinics in sports medicine. 2011 Jan 1;30(1):33–48.

9 Gallo V, Motley K, Kemp SP, Mian S, Patel T, James L, Pearce N, McElvenny D. Concussion and long-term cognitive impairment among professional or elite sport-persons: a systematic review. Journal of Neurology, Neurosurgery & Psychiatry. 2020 May 1;91(5):455–68.

10 Cook MJ, Gardner AJ, Wojtowicz M, Williams WH, Iverson GL, Stanwell P. Task-related functional magnetic resonance imaging activations in patients with acute and subacute mild traumatic brain injury: A coordinate-based meta-analysis. Neuroimage: clinical. 2020 Jan 1;25:102129.

11 Tan SJ, Kerr G, Sullivan JP, Peake JM. A brief review of the application of neuroergonomics in skilled cognition during expert sports performance. Frontiers in human neuroscience. 2019;13:278.

12 Wise SP, Di Pellegrino G, Boussaoud D. The premotor cortex and nonstandard sensorimotor mapping. Canadian journal of physiology and pharmacology. 1996 Apr 1;74(4):469–82.

13 Granek JA, Sergio LE. Evidence for distinct brain networks in the control of rule-based motor behavior. Journal of neurophysiology. 2015 Aug;114(2):1298–309.

14 Dalecki M, Albines D, Macpherson A, Sergio LE. Prolonged cognitive–motor impairments in children and adolescents with a history of concussion. Concussion. 2016 May 12;1(3):CNC14.

15 Dalecki M, Usand J, Van Gemmert AW, Sergio LE. Motor deficits in youth with concussion history: issues with task novelty or task demand?. International journal of sports medicine. 2020 Sep 1;41(10):688–95.

16 Brown JA, Dalecki M, Hughes C, Macpherson AK, Sergio LE. Cognitive-motor integration deficits in young adult athletes following concussion. BMC sports science, medicine and rehabilitation. 2015 Dec 1;7(1):25.

17 Hurtubise J, Gorbet D, Hamandi Y, Macpherson A, Sergio L. The effect of concussion history on cognitive-motor integration in elite hockey players. Concussion. 2016 Sep 6;1(3):CNC17.

18 Gorbet DJ, Sergio LE. Don’t watch where you’re going: The neural correlates of decoupling eye and arm movements. Behavioural brain research. 2016 Feb 1;298:229–40.

19 Gorbet DJ, Sergio LE. Move faster, think later: Women who play action video games have quicker visually-guided responses with later onset visuomotor-related brain activity. PloS one. 2018 Jan 24;13(1):e0189110.

20 Granek JA, Gorbet DJ, Sergio LE. Extensive video-game experience alters cortical networks for complex visuomotor transformations. Cortex. 2010 Oct 1;46(9):1165–77.

21 Lotze M, Scheler G, Tan HR, Braun C, Birbaumer N. The musician’s brain: functional imaging of amateurs and professionals during performance and imagery. Neuroimage. 2003 Nov 1;20(3):1817–29.

22 Haslinger B, Erhard P, Altenmüller E, Hennenlotter A, Schwaiger M, Gräfin von Einsiedel H, Rummeny E, Conrad B, Ceballos-Baumann AO. Reduced recruitment of motor association areas during bimanual coordination in concert pianists. Human brain mapping. 2004 Jul;22(3):206–15.

23 Jäncke L, Shah NJ, Peters M. Cortical activations in primary and secondary motor areas for complex bimanual movements in professional pianists. Cognitive Brain Research. 2000 Sep 1;10(1-2):177–83.

24 Krings T, Töpper R, Foltys H, Erberich S, Sparing R, Willmes K, Thron A. Cortical activation patterns during complex motor tasks in piano players and control subjects. A functional magnetic resonance imaging study. Neuroscience letters. 2000 Jan 14;278(3):189–93.

25 Petrini K, Pollick FE, Dahl S, McAleer P, McKay L, Rocchesso D, Waadeland CH, Love S, Avanzini F, Puce A. Action expertise reduces brain activity for audiovisual matching actions: an fMRI study with expert drummers. Neuroimage. 2011 Jun 1;56(3):1480–92.

26 Dalecki M, Gorbet DJ, Macpherson A, Sergio LE. Sport experience is correlated with complex motor skill recovery in youth following concussion. European journal of sport science. 2019 Oct 21;19(9):1257–66.

27 Gorbet DJ, Sergio LE. Preliminary sex differences in human cortical BOLD fMRI activity during the preparation of increasingly complex visually guided movements. European Journal of Neuroscience. 2007 Feb;25(4):1228–39.

28 Li X, Morgan PS, Ashburner J, Smith J, Rorden C. The first step for neuroimaging data analysis: DICOM to NIfTI conversion. Journal of neuroscience methods. 2016 May 1;264:47–56.

29 Kundu P, Inati SJ, Evans JW, Luh WM, Bandettini PA. Differentiating BOLD and non-BOLD signals in fMRI time series using multi-echo EPI. Neuroimage. 2012 Apr 15;60(3):1759–70.

30 Beckmann CF, Smith SM. Probabilistic independent component analysis for functional magnetic resonance imaging. IEEE transactions on medical imaging. 2004 Feb 6;23(2):137–52.

31 Yeo BT, Krienen FM, Sepulcre J, Sabuncu MR, Lashkari D, Hollinshead M, Roffman JL, Smoller JW, Zöllei L, Polimeni JR, Fischl B. The organization of the human cerebral cortex estimated by intrinsic functional connectivity. Journal of neurophysiology. 2011 Sep 1.

32 Bernard JA, Seidler RD, Hassevoort KM, Benson BL, Welsh RC, Wiggins JL, Jaeggi SM, Buschkuehl M, Monk CS, Jonides J, Peltier SJ. Resting state cortico-cerebellar functional connectivity networks: a comparison of anatomical and self-organizing map approaches. Frontiers in neuroanatomy. 2012 Aug 10;6:31.

33 Wright DK, Gardner AJ, Wojtowicz M, Iverson GL, O’Brien TJ, Shultz SR, Stanwell P. White Matter Abnormalities in Retired Professional Rugby League Players with a History of Concussion. Journal of Neurotrauma. 2020 Jun 3.

34 Nickerson LD, Smith SM, Öngür D, Beckmann CF. Using dual regression to investigate network shape and amplitude in functional connectivity analyses. Frontiers in neuroscience. 2017 Mar 13;11:115.

35 Churchill NW, Caverzasi E, Graham SJ, Hutchison MG, Schweizer TA. White matter microstructure in athletes with a history of concussion: comparing diffusion tensor imaging (DTI) and neurite orientation dispersion and density imaging (NODDI). Human brain mapping. 2017 Aug;38(8):4201–11.

36 Cook MJ, Gardner AJ, Wojtowicz M, Williams WH, Iverson GL, Stanwell P. Task-related functional magnetic resonance imaging activations in patients with acute and subacute mild traumatic brain injury: A coordinate-based meta-analysis. Neuroimage: clinical. 2020 Jan 1;25:102129.

37 Hristopulos DT, Babul A, Shazia’Ayn Babul LR, Virji-Babul N. Disrupted information flow in resting-state in adolescents with sports related concussion. Frontiers in Human Neuroscience. 2019;13.

38 Czerniak SM, Sikoglu EM, Navarro AA, McCafferty J, Eisenstock J, Stevenson JH, King JA, Moore CM. A resting state functional magnetic resonance imaging study of concussion in collegiate athletes. Brain imaging and behavior. 2015 Jun 1;9(2):323–32.

39 Cubon VA, Murugavel M, Holmes KW, Dettwiler A. Preliminary evidence from a prospective DTI study suggests a posterior-to-anterior pattern of recovery in college athletes with sports-related concussion. Brain and behavior. 2018 Dec;8(12):e01165.

40 Murdaugh DL, King TZ, Sun B, Jones RA, Ono KE, Reisner A, Burns TG. Longitudinal changes in resting state connectivity and white matter integrity in adolescents with sports-related concussion. Journal of the International Neuropsychological Society. 2018 Sep;24(8):781–92.

41 Bazarian JJ, Blyth B, Mookerjee S, He H, McDermott MP. Sex differences in outcome after mild traumatic brain injury. Journal of neurotrauma. 2010 Mar 1;27(3):527–39.

42 Ono KE, Burns TG, Bearden DJ, McManus SM, King H, Reisner A. Sex-based differences as a predictor of recovery trajectories in young athletes after a sports-related concussion. The American journal of sports medicine. 2016 Mar;44(3):748–52.

43 Streifer M, Brown AM, Porfido T, Anderson EZ, Buckman JF, Esopenko C. The potential role of the cervical spine in sports-related concussion: Clinical perspectives and considerations for risk reduction. journal of orthopaedic & sports physical therapy. 2019 Mar;49(3):202–8.

44 Gallagher V, Kramer N, Abbott K, Alexander J, Breiter H, Herrold A, Lindley T, Mjaanes J, Reilly J. The effects of sex differences and hormonal contraception on outcomes after collegiate sports-related concussion. Journal of Neurotrauma. 2018 Jun 1;35(11):1242–7.

45 Hamer J, Churchill NW, Hutchison MG, Graham SJ, Schweizer TA. Sex differences in cerebral blood flow associated with a history of concussion. Journal of Neurotrauma. 2020 May 15;37(10):1197–203.

46 Dumais KM, Chernyak S, Nickerson LD, Janes AC. Sex differences in default mode and dorsal attention network engagement. PLoS One. 2018 Jun 14;13(6):e0199049.

47 McCorkle TA, Giacometti LL, Khurana S, Raghupathi R. Sex differences in cognitive deficits following repetitive mild TBI in adolescent rats. Program No. 300.11. 2019 Neuroscience Meeting Planner. Chicago, IL: Society for Neuroscience, 2019

48 Palmer SJ, Ng B, Abugharbieh R, Eigenraam L, McKeown MJ. Motor reserve and novel area recruitment: amplitude and spatial characteristics of compensation in Parkinson’s disease. European Journal of Neuroscience. 2009 Jun;29(11):2187–96.

49 Mortimer JA, Schuman LM, French LR. Epidemiology of dementing illness. The epidemiology of dementia: Monographs in epidemiology and biostatistics. 1981:323–33.

50 Satz P. Brain reserve capacity on symptom onset after brain injury: a formulation and review of evidence for threshold theory. Neuropsychology. 1993 Jul;7(3):273.

51 Olazaran J, Muñiz R, Reisberg B, Peña-Casanova J, Del Ser T, Cruz-Jentoft AJ, Serrano P, Navarro E, de la Rocha MG, Frank A, Galiano M. Benefits of cognitive-motor intervention in MCI and mild to moderate Alzheimer disease. Neurology. 2004 Dec 28;63(12):2348–53.

52 Kandel S, Orliaguet JP, Boe LJ. Detecting anticipatory events in handwriting movements. Perception. 2000 Aug;29(8):953–64.

53 Gorbet DJ, Sergio LE. The behavioural consequences of dissociating the spatial directions of eye and arm movements. Brain research. 2009 Aug 11;1284:77–88.

54 Miall RC, Reckess GZ, Imamizu H. The cerebellum coordinates eye and hand tracking movements. Nature neuroscience. 2001 Jun;4(6):638–44.

55 Teel EF, Marshall SW, Shankar V, McCrea M, Guskiewicz KM. Predicting recovery patterns after sport-related concussion. Journal of athletic training. 2017 Mar;52(3):288–98.

56 Broglio SP, Sosnoff JJ, Ferrara MS. The relationship of athlete-reported concussion symptoms and objective measures of neurocognitive function and postural control. Clinical Journal of Sport Medicine. 2009 Sep 1;19(5):377–82.

57 Sabes PN. The planning and control of reaching movements. Current opinion in neurobiology. 2000 Dec 1;10(6):740–6.

58 Wolpert, D., Ghahramani, Z., & Jordan, M. (1995). An internal model for sensorimotor integration. Science, 269(5232), 1880–1882.

59 Hawkins, K., Goyal, A., & Sergio, L. (2015). Diffusion tensor imaging correlates of cognitive-motor decline in normal aging and increased Alzheimer’s disease risk. Journal of Alzheimer’s disease, 44(3), 867–878

60 Hurtubise, J. M., Gorbet, D. J., Hynes, L. M., Macpherson, A. K., & Sergio, L. E. (2020). White matter integrity and its relationship to cognitive-motor integration in females with and without post-concussion syndrome. Journal of Neurotrauma.

